# Leveraging Experimental Vasculature Data for High Resolution Brain Tumor Simulations

**DOI:** 10.1101/2024.11.25.625136

**Authors:** Eric Behle, Julian Herold, Alexander Schug

## Abstract

Cancer remains a leading cause of mortality. Multidisciplinary studies probe its complex pathology to increase treatment options. Computational modeling of tumor growth on high-performance computing resources offers microscopic insight into its progress and a valuable avenue for advancing our understanding. However, the effective initialization and parameterization of the underlying models require high-resolution data from real tissue structures. Here, we leveraged high performance computing resources and a massive dataset of a mouse brain’s entire vascular network. We processed these image stacks into detailed three-dimensional representations, identified brain regions of interest, and conducted a series of large-scale simulations to investigate how tumor growth is influenced by local vascular network characteristics. By simulating tumor growth with sub-cellular resolution, we can probe to which extent vessel density and vessel network length influence tumor growth. We determined that vessel density is the primary determinant of growth rate. Finally, our results allowed us to extrapolate tumor cell growth predictions for the entire mouse brain, highlighting the critical role of vascular topology in tumor progression. Such increasingly realistic simulations of cancer cells and their microenvironment enable researchers to increasingly bridge the gap between basic biology and clinical practice, ultimately supporting the development of more effective personalized cancer therapies.

## 1 Introduction

Cancer remains one of the leading causes of death worldwide. Thus, it poses an ever-present global health challenge that urgently requires improved therapeutic strategies and deeper insights into its pathology. Despite remarkable advances in cancer treatment, the complexity of tumor growth, its interaction with surrounding tissues, and its eventual resistance to therapies often lead to challenges in effective clinical management. Central to addressing these challenges are advanced simulations of cancer tissue. The massive growth of compute resources allows studies which offer the possibility of highly detailed, sub-cellular insights into the dynamics driving tumor progression. They also enable researchers to visualize and analyze critical interactions within the tumor microenvironment, that are otherwise challenging to study in vivo. Such interactions include nutrient gradients, vascular influences, and cellular displacement. By mimicking these intricate biological processes in a controlled computational setting, simulations have emerged as a vital pillar in cancer research. They bridge the gap between empirical experimental data and theoretical models, complementing wet-lab findings and aiding in hypothesis testing. [2–5]. Computational simulations have been instrumental not only in cancer studies but also in fields such as developmental biology, where they provide insights into tissue mechanics and cellular processes in embryogenesis and wound healing. This multidisciplinary approach facilitates the development of comprehensive models that can inform various stages of cancer development and therapeutic response. These can vary in resolution and scope, from modeling cellular and molecular dynamics at a fine scale to examining tissue-wide phenomena [6, 7]. The ultimate goal of many research initiatives is to develop “digital twins” of tumors: highly realistic, computational representations of tumors within healthy tissue that mimic their real-life counterparts [8–10]. These digital twins promise to revolutionize personalized medicine, enabling simulations of disease progression and treatment response specific to an individual’s unique biological profile. Building such a model requires an accurate and dynamic morphology of both the healthy and tumorous tissue. It needs to capture the growth behavior of tumors as they interact with the cellular environment, migrate, and displace other cells. A key factor in these simulations is nutrient availability, as nutrient gradients largely dictate tumor growth dynamics and progression patterns [11, 12]. The transition from avascular (limited to nutrient diffusion) to vascularized growth phases is a significant turning point in tumor development, where angiogenesis, the formation of new blood vessels, provides the tumor with an essential nutrient supply. Without this transition, tumors cannot grow beyond a critical volume of around 1-3 mm^3^ before nutrient limitations halt further expansion. Therefore, the study of nutrient supply mechanisms and vascularization processes is a focal point in cancer research, with numerous studies investigating how nutrient gradients impact tumor viability and aggressiveness[13–15]. Previous simulations relied on idealized representations of nutrient distributions, such as simple radial gradients that assume uniformity in nutrient availability[16]. However, these models lack the complexity of real tissue environments, where nutrient distribution is influenced by an intricate vascular network. A more realistic depiction of the tissue microenvironment requires detailed information about vascular topology, particularly at the capillary level, which is essential for understanding how tumors respond to spatially heterogeneous nutrient supplies. Here, we aim to go beyond idealized models by integrating data that closely approximates actual tissue environments. Leveraging recent advances in fluorescent microscopy and image processing, we utilize high-resolution data from Di Giovanna et al. [1], which provides a capillary-scale map of the entire vascular network within a mouse brain. Such data allow us to incorporate highly realistic tissue morphology into our simulations, capturing the nuanced interactions between tumor cells and their vascular surroundings. Our approach takes advantage of computational resources and simulation frameworks. High-performance computing clusters, now more accessible than ever, support these complex models by handling vast datasets and performing simulations at sub-cellular resolution. A key factor in executing these simulations is the parallelization and scalability of simulation software; the CiS (Cells in Silico) framework [17], used in our study, demonstrates effective performance scaling across thousands of computing nodes, making it a suitable tool for handling the demands of high-resolution, three-dimensional simulations.

In the following, we will first outline the dataset, describing the imaging techniques and data processing methods used to achieve a detailed three-dimensional representation of vascular topology. We will then discuss the integration of these vascular maps into our simulation pipeline and present our analysis of how variations in vascular network characteristics influence tumor growth patterns. A critical question in tumor growth is the dependence of growth on the local microenvironment and accessibility of nutrients. Here, large scale simulations provide sub-cellular insight and identify key factors catalyzing growth. Finally, we will summarize the insights gained from these simulations and propose directions for future research, including potential applications in the development of patient-specific cancer therapies.

## 2 Results

### 2.1 Vascular Data Processing for Large-Scale Simulations

To inform our simulations based on the vascular network topology, we first processed the raw data into analyzable structures. Here, we detail this processing pipeline.

#### 2.1.1 Raw Microscopy data

Di Giovanna et al. measured the entire vascular network of a mouse brain at capillary resolution [1] (see Figure 1 and 2). The raw data generated by them are available in the form of z-stacks of light sheet microscopy images [1]. Each stack contains 2160 microscopy images with a resolution of 2048 × 2048 px. One px represents an area of 0.65 × 0.65 µm^2^, and the spacing between each image is 2 µm in z-direction. Hence, a single stack depicts a volume of 1331 × 1331 × 4320 µm^3^. Furthermore, adjacent stacks have an overlap of roughly 300 µm in x- and y-direction.

**Fig. 1.**
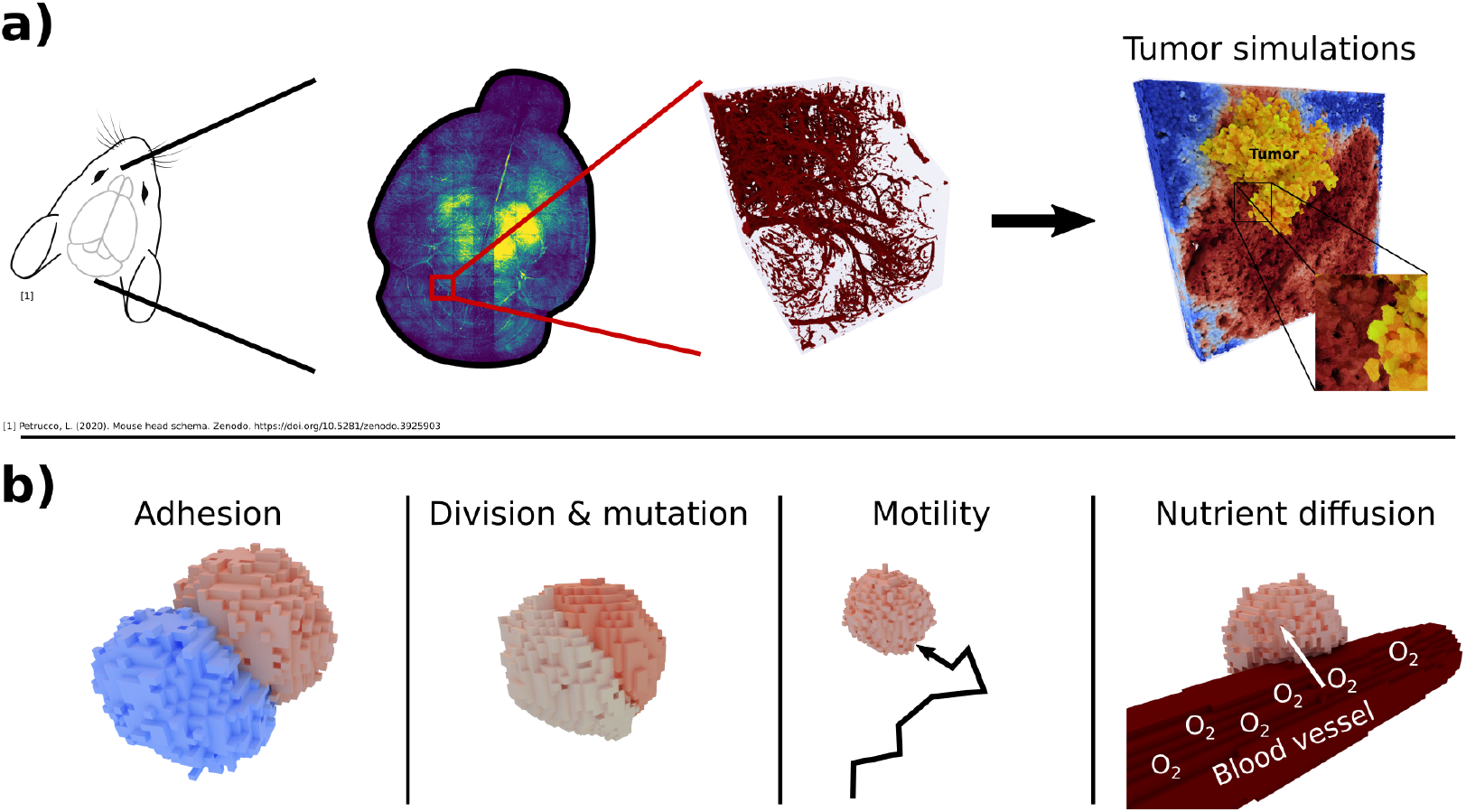
Visual abstract. Mouse brain vasculature microscopy data are a useful resource for the initialization of tumor growth simulations. We utilized data by Di Giovanna et al. [1], who imaged the vasculature network of an entire mouse brain and provided light-sheet microscopy z-stacks. In this study, we implemented a pipeline to clean, align and combine these stacks, obtaining three-dimensional representations. We then performed an analysis of the network topology at different positions and used this analysis to identify areas of interest within the brain. We then performed a large scale analysis of tumor growth simulations depending on the nutrient environment based on these regions of interest.

**Fig. 2.**
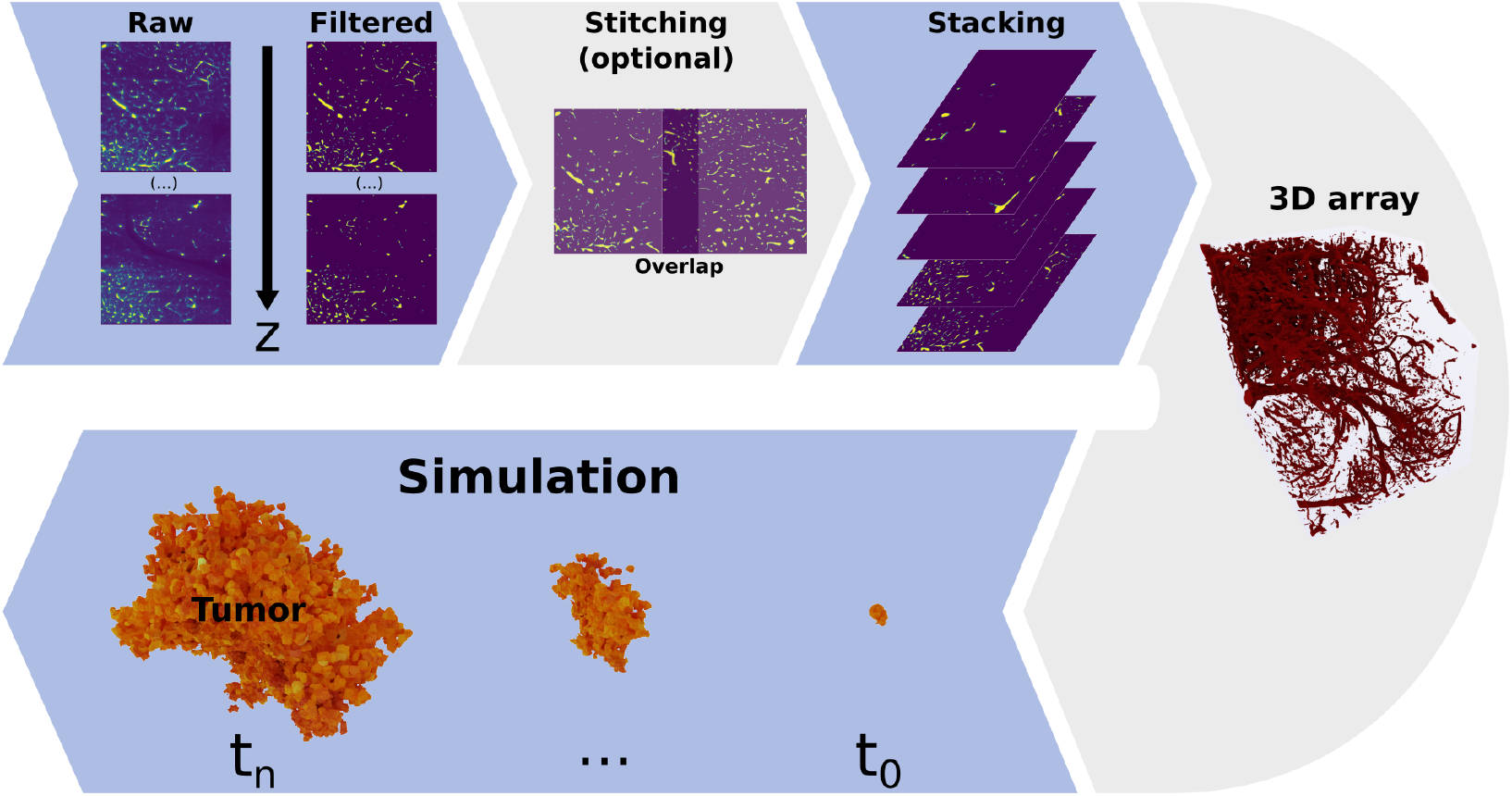
Overview of the mouse brain vasculature data processing pipeline. The pipeline begins with raw microscopy images, which are converted into binary masks using a threshold filter to isolate vascular structures. Due to the size limitation of individual z-stacks (1331 *×* 1331 *×* 4320 µm^3^), a stitching procedure can be applied to extend the simulation volume. By optimizing the alignment of adjacent z-stacks, we minimize artifacts in the merged image. These binary masks are then assembled into a 3D array, which is subsequently imported into the *Cells in Silico* (CiS) platform, providing spatial and structural information for nutrient source modeling in our simulations.

#### 2.1.2 Processing into 3D volumes

Processing the data into a usable form presents several challenges. First, since they are raw microscopy images, they contain noise. Secondly, there may be artifacts from the experimental procedure, especially at the edges of each stack. Therefore, the first step in obtaining a three-dimensional representation of all stacks was to denoise them. Towards this goal, we built binary masks from the microscopy images of each of them. This was done by applying a threshold filter on each image. All images were then combined in a three-dimensional array. The resulting array contains the value 1 in locations where a blood vessel was present, and the value 0 otherwise. Next, we rescaled this three-dimensional binary mask. By doing so we matched the native resolution of our model, such that 1 voxel represents a volume of 1 µm^3^. This was done for each individual microscopy stack. Finally, in order to obtain a roughly cubic geometry for our simulations, we split each stack into four substacks along the z-axis. The final volume of the binary mask of each substack was therefore *V* = 1331×1331×1080 µm^3^.

#### 2.1.3 Stitching multiple stacks

We implemented a method that enables us to stitch neighboring regions into a single stack. This addresses the second challenge of the raw data by minimizing artifacts at the stack edges. See SI appendix A for more details.

#### 2.1.4 Microscopy stack grouping

Even though supercomputers have grown immensely in computational power in recent years [18], simulating the entire 3D representation of the mouse brain at µm-resolution would still be a monumental challenge. On the other hand, it could be argued that due to the respective similarity between different regions (see also SI Figure 5), significant insights can already be gained by focusing on a subset of archetypes. To determine these archetypes, we decided to characterize the substacks and find points of interest. We used two parameters for this characterization: the vessel network density *ρ*, and a parameter which we call the network length fraction *f*_l_. *ρ* was taken as the fraction of vessel voxels within the substack. *f*_l_ was calculated through a skeletonizing procedure [19]. Here, each binary mask was eroded, reducing the diameter of each vessel to 1 voxel. *f*_l_ is obtained by computing the fraction of remaining to originally present vessel voxels. A value of *f*_l_ close to 0 indicates a low number of thick blood vessels and hence a low overall network length. A value close to 1 indicates many thin vessels which increases the overall network length. *f*_l_ can also be viewed as a measure of the relative surface area of the vessel network. As seen in Figure 3 a), k-means clustering by *ρ* and *f*_l_ does not yield fully distinct clusters, except for B and C. Nonetheless, it enables us to choose groups of interest much better than randomly picking from all substacks. For the following simulation study, we chose the substacks which were closest to the cluster centroids (red points in Figure 3 a)). A snapshot of each chosen stack’s three-dimensional structure is also shown in Figure 3 b).

**Fig. 3.**
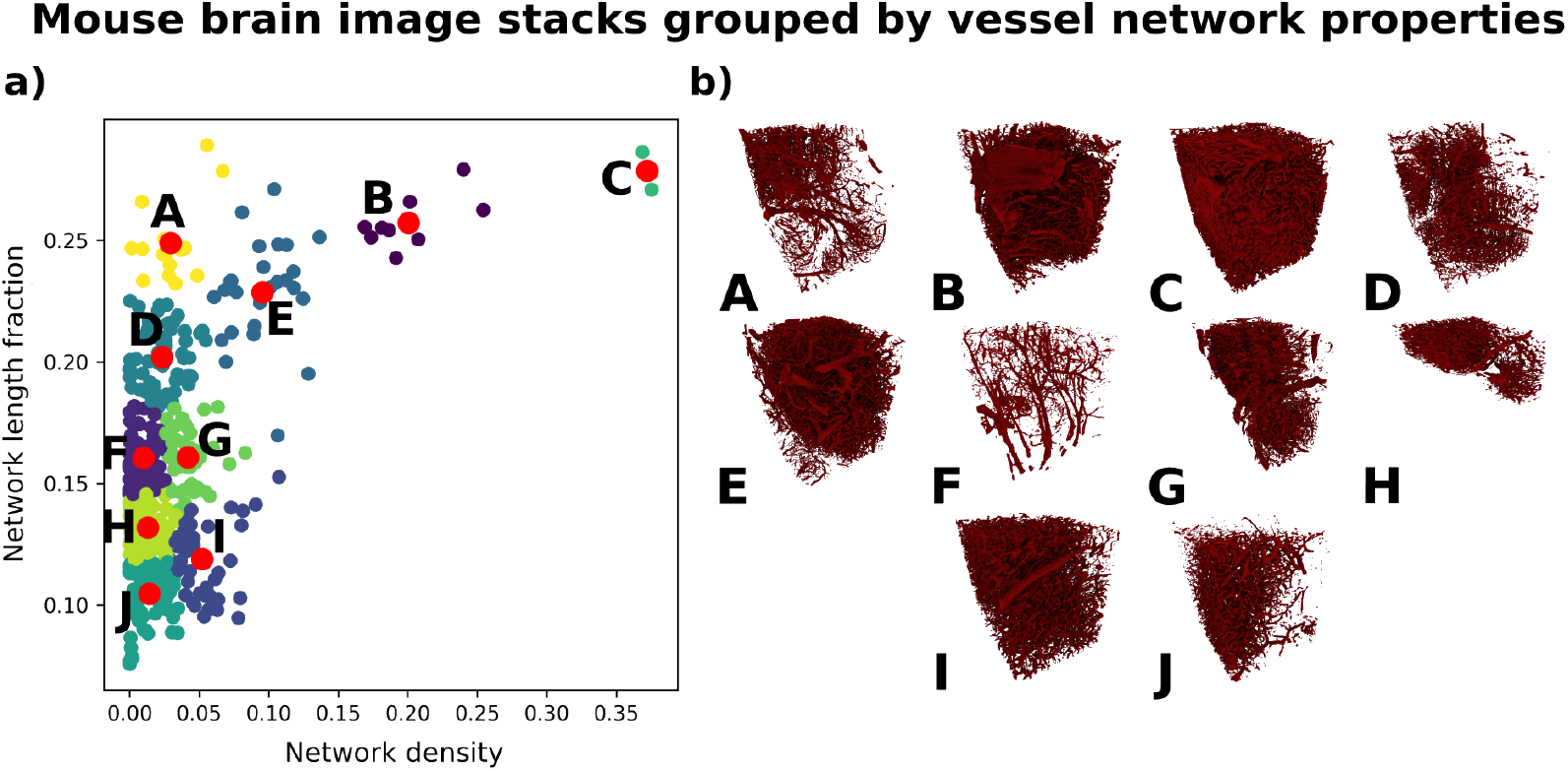
Detection of points of interest within mouse brain vasculature data. **a)** Processed mouse brain microscopy stacks arranged regarding their local blood vessel network properties. Shown are, for each stack, the network density *ρ*, and the network length fraction *f*_l_. The color indicates the cluster membership obtained from kmeans-clustering. The red points represent the centroid of each cluster. **b)** Snapshot of the three-dimensional structure of the stacks chosen for further study. Each chosen stack is the one closest to the centroid of each cluster.

### 2.2 Influence of vessel network topology on tumor growth

After choosing areas of interest within the mouse brain, we investigated the influence of *ρ* and *f*_l_ on tumor growth. Here, we wanted to test two non-exclusive hypotheses regarding nutrient driven tumor growth:

- Tumor growth is mostly driven by the density of the vascular network (*ρ*). In this case the local blood vessels drive tumor growth, since higher density leads to more available nutrients.
- Tumor growth is mostly driven by the local topology of the vascular network (*f*_l_). In this case different local topologies result in differences of the vessel network surface area, influencing the available amount of nutrients.

The simplest assumption was that there is a linear relationship between *ρ* and the final tumor cell count. To evaluate this, we performed simulations of tumors seeded in different parts of the brain using the *Cells in Silico* (CiS) tissue simulation framework [17]. See section 4.1 for a detailed summary of CiS.

#### 2.2.1 Simulations

In each simulation, blood vessels were represented by inserting specialized voxels into the Cellular Potts Model (CPM) layer of the CiS framework (see 4.3). These vessel voxels, derived from the binary mask data from section 2.1.2, served as nutrient sources. The remaining volume was populated with healthy tissue cells, maintaining a density consistent with mouse brain cell populations (∼130,000 cells) [20]. A single initial tumor, consisting of approximately 10 cells, was then seeded.

Nutrient diffusion was modeled using the CiS compound diffusion layer (see section 4.1), with a single nutrient acting as a proxy for oxygen, glucose, and other essential compounds delivered by blood vessels. Tumor cells could proliferate if they met the necessary nutrient threshold. With each division, a tumor cell could undergo a mutation affecting either its cell-cell adhesion strength or its motility. Additionally, each cell consumed nutrients, and tumor cells would die if their nutrient levels dropped below a critical threshold.

It is important to note that the traditional CPM, as used here, does not fully capture the intricate morphologies of neurons and glial cells. Nonetheless, CPMs are widely applied in similar studies for tissue modeling [21, 22]. We argue that this simplification is justified, as the primary cell-cell interactions driving these simulations — adhesion forces and nutrient diffusion — are closely related to the shared surface area between cells. Introducing more complex cell shapes would mainly result in two effects: (a) increasing shared surface area for cells already in contact, effectively modifying adhesion and diffusion parameters for all cells uniformly, and (b) creating small shared surfaces between cells previously not in contact. Since the first effect would scale uniformly and the second effect introduces minimal interaction, the impact of these adjustments would be limited. Moreover, adding complex shapes would significantly increase both model and computational complexity, with only marginal gains in accuracy.

More details on the model parameters are provided in section 4.2 and in SI table 1. We performed eight simulations for each centroid substack obtained in section 2.1.4. In each simulation, the tumor was seeded at one of eight points. These points were at one third and two thirds of each simulation box diagonal. Each simulation ran for an total amount of 250 000 MC steps. A snapshot of a single simulation after 100 000 Monte Carlo (MC) steps is provided in Figure 4.

**Fig. 4.**
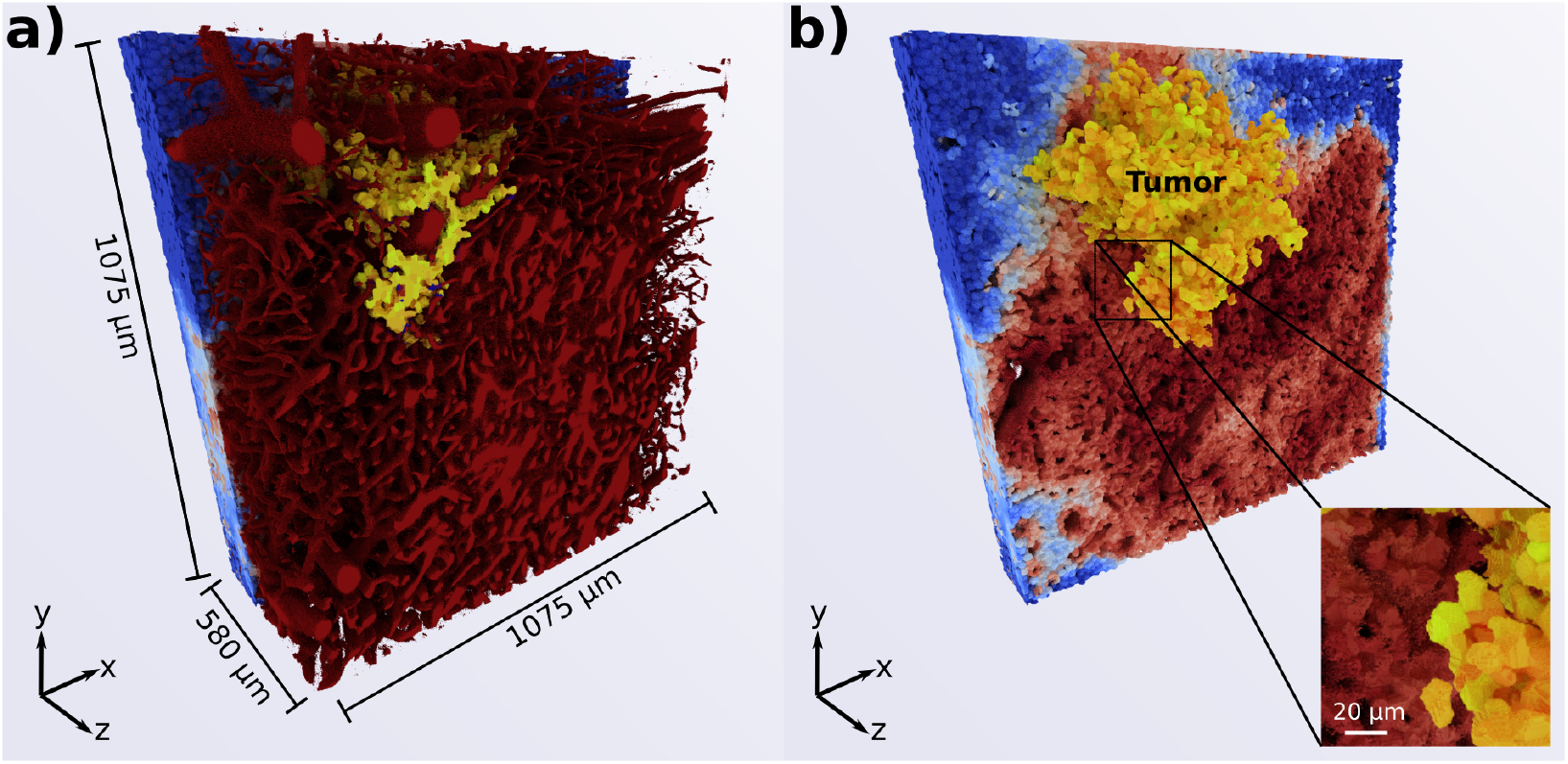
Visualization of a simulated tumor growing within a mouse brain vasculature environment (cluster B in Fig. 3b) after 100,000 Monte Carlo (MC) steps. **(a)** The dark red structure represents the blood vessel network, providing nutrient sources. **(b)** The tumor cells (yellow) are shown expanding in three dimensions, with the surrounding healthy cells color-coded according to their nutrient content (blue to red gradient), indicating the spatial distribution of nutrient availability. The displayed volume is a subset of the entire simulation space (1344 *×* 1344 *×* 1080 µm^3^) to focus on the tumor’s immediate environment.

**Fig. 5.**
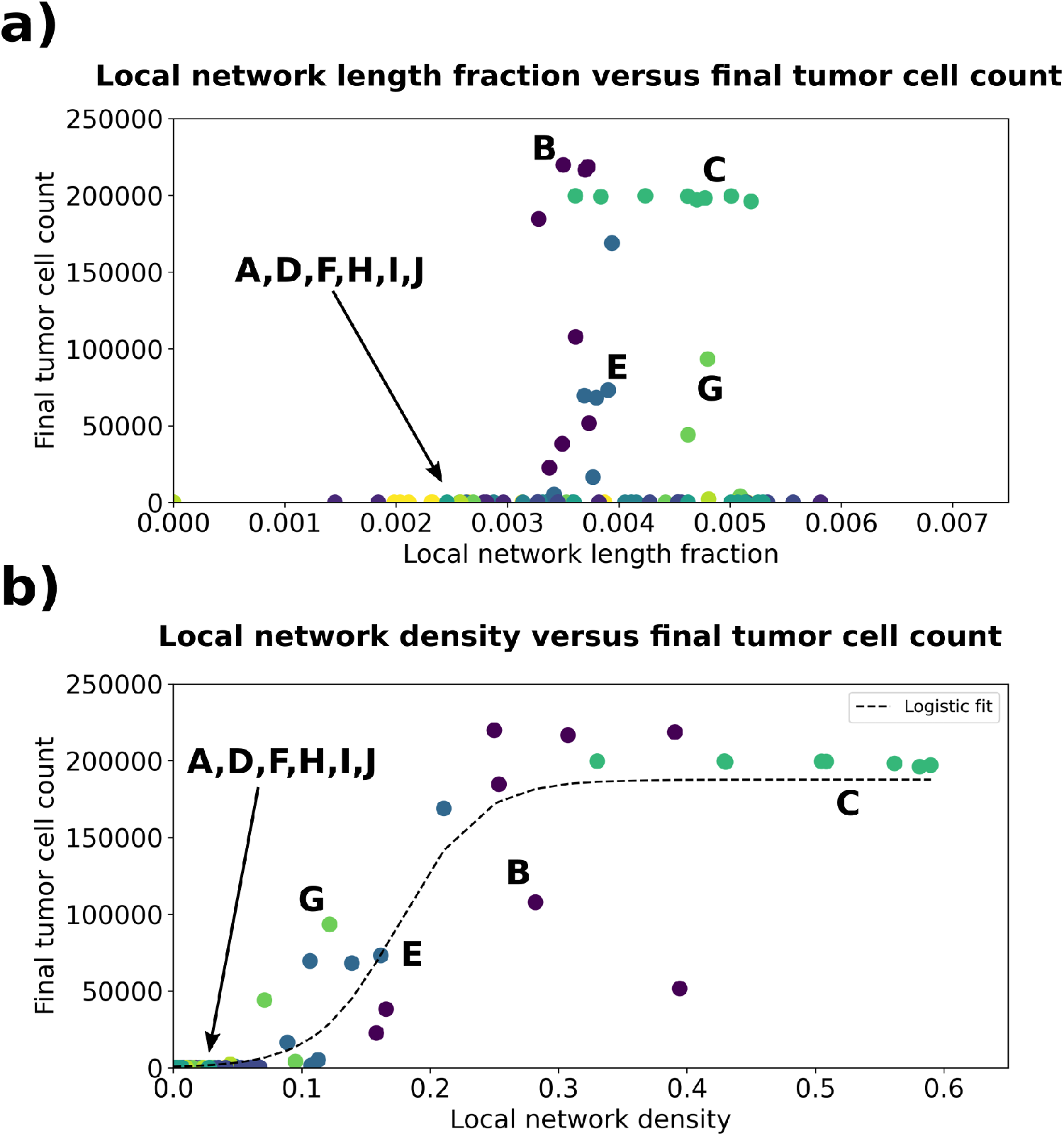
Mousebrain vasculature property comparison **a)** Network length fraction versus final tumor cell count. Shown are the color-coded final tumor cell counts for eight simulations of each respective microscopy stack chosen in section 2.1.4. In each simulation, the tumor was placed at a different starting position, and hence the local network length fraction differs. The network length fraction serves as a proxy for the vessel network length. We see no clear dependence of the final tumor cell count on the network length. **b)** Local blood vessel density versus final tumor cell count. Shown are the color-coded final tumor cell count for eight simulations of each respective microscopy stack chosen in section 2.1.4. In each simulation, the tumor was placed at a different starting position, and hence the local blood vessel density differs. The growth behavior follows a logistic curve.

#### 2.2.2 Tumor growth behavior

To compare all simulations, we first calculated the local blood vessel properties in the vicinity of each tumor. For this we chose a sphere of radius 250 µm around the initial tumor center. We then calculated the local density *ρ*_local_ and local network length fraction *f*_l,local_ within this sphere. For the calculation of *f*_l,local_ we had to further refine our procedure, since we noted artifacts which distorted the skeletonization. After applying a post-processing pipeline of morphological operations, this was improved. Finally, we determined the number of tumor cells at the end of each simulation and compared the quantities in Figure 5. While the relationship between the network length fraction and the final tumor cell count is less clear, the dependence of the final tumor cell count on the local blood vessel density shown in Figure 5 b) roughly follows a logistic curve:

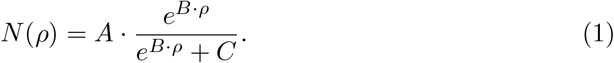

These data show that the simple assumption of a linear dependence between vessel density and tumor growth rate does not hold. Instead there exists a threshold above which the tumor grows rapidly. Furthermore, the higher blood vessel surface area resulting from the high network length fractions does not appear to be a main driver for tumor growth.

#### 2.2.3 Predicted tumor growth in the entire mouse brain

We can use the relation between local blood vessel density and final tumor cell count to estimate the tumor size for the entire mouse brain by fitting equation 1 to the data shown in Figure 5 b) (see also SI table 2). We used the resulting values to calculate the expected cell count for the given vessel density *ρ* of each substack. Within our simulation environment tumors are expected to grow largest in the densest regions in the center of the brain, and to not grow well in the periphery (see Figure 6 and SI Figure 3).

**Fig. 6.**
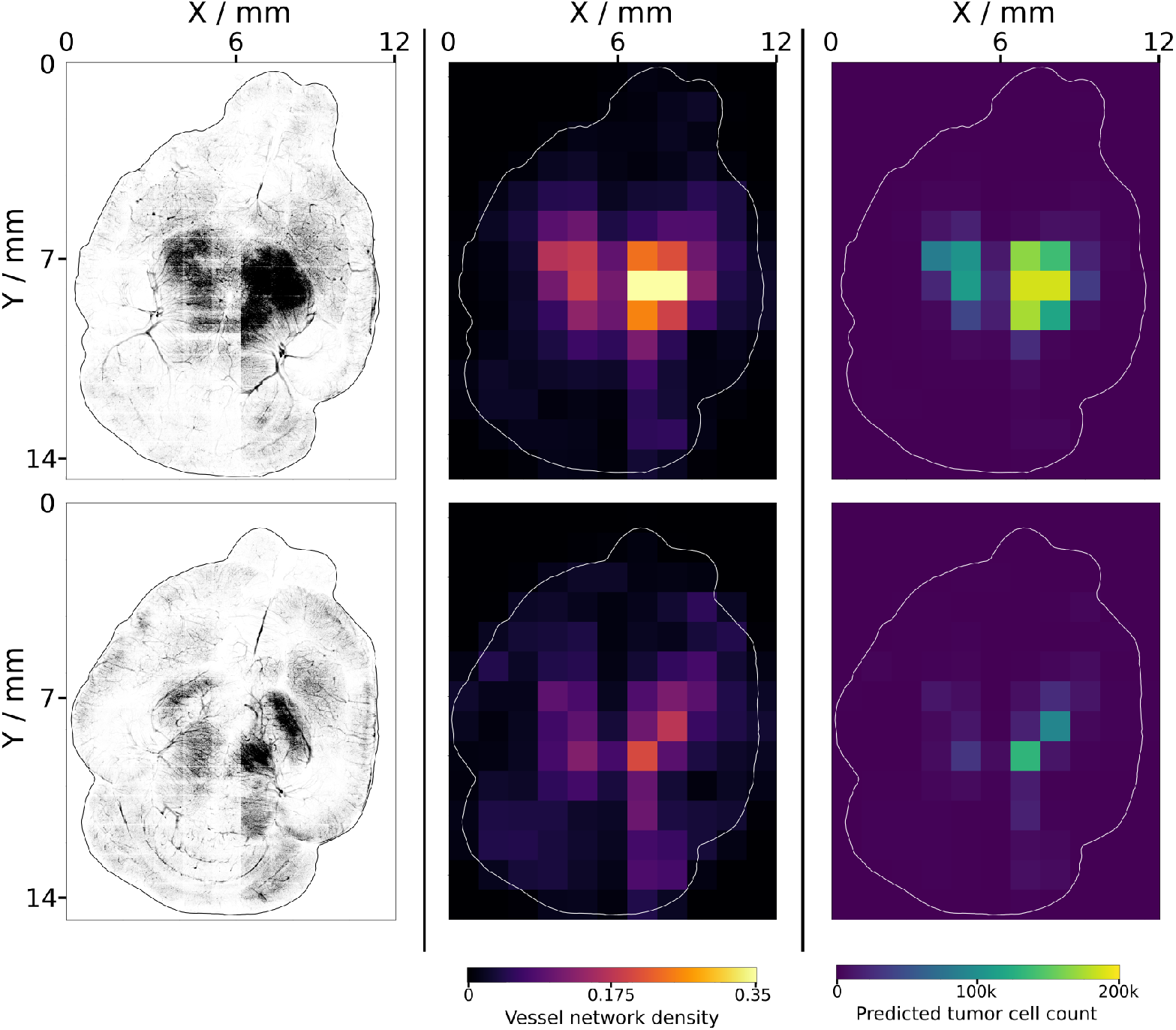
Mouse brain blood vessel density and predicted tumor growth. Left: Top-down views of the processed and stitched microscopy stacks for the lowest two layers of the brain. Middle: heatmaps showing the blood vessel volume fraction for each microscopy stack. Right: heatmaps showing the predicted tumor cell count for each microscopy stack. Predicted values were obtained using equation 1 with the values from fitting to the simulation data (see section 2.2.3 and SI table 2.

## 3 Discussion and conclusion

Computational studies of tumor growth can profit immensely from existing experimental data. The challenge is to incorporate these data in a meaningful manner. In this study, we showed how we utilized highly detailed microscopy data on mouse brain vasculature in order to inform our simulation framework CiS. After developing a data processing pipeline for the microscopy stacks found in the raw data, we characterized each processed stack by its vessel network topology. We utilized two parameters for this characterization: the blood vessel density *ρ*, and the network length fraction *f*_l_. After choosing ten substacks of interest based on the two parameters, we performed simulations of tumor growth in this wide range of nutrient environments. Placing the initial tumor in different locations yielded different local starting topologies, and we observed different growth behavior based on this. We saw that the main driver of tumor growth is the local vessel density, and that the final cell count roughly follows a logistic curve. While there are some outliers in Figure 5 b), these show that the final tumor cell count is not determined solely by the starting position of the tumor, but also its surrounding further from it (see also SI Figure 4).

The network length fraction, which approximates the distribution of vessel thicknesses, does not appear to have significant influence on the final tumor size. This was surprising to us, since we expected that a higher network length fraction would correspond to a larger surface area of the vessels, leading to increased nutrient diffusion.

Using a logistic curve fitted to the local density data, we predicted the tumor growth in the entire brain. Strong tumor growth was predicted mainly in the dense regions in the center of the lower two planes of the brain. One major caveat to mention here is that we used the global vessel density per stack for the prediction, and did not look at more localized vessel properties. This effectively represents an averaging of larger volumes, thus losing information about inhomogeneities. Hence, we might miss some regions which locally surpass the established threshold. Furthermore, as seen in the left pane of Figure 6, there are some residual brightness artifacts from the microscopy, which may influence the results.

This study shows that there still is a large amount of information to be gained from the existing experimental data on tumors. We aim to utilize these data for further studies of more realistic tumors. In particular, the stitching procedure that we mentioned in section 2.1 could be used to simulate tumors of multiple mm^3^ in volume. Such system sizes would enable us to perform *in silico* radio- and chemotherapy treatment studies in the future. Furthermore, it is a step on the way to simulating fully vascularized macroscopic tumor growth with our model. Our future studies will focus on implementing tumor-induced angiogenesis, the remaining step on this path.

## 4 Materials and Methods

### 4.1 Model description

*Cells in Silico* was developed by our group as a framework for simulating the dynamics of cells and tissues at subcellular resolution [16, 17, 23]. It is a hybrid model composed of a Cellular Potts Model (CPM), nutrient and signal exchange functionalities, and an agent-based layer. With it, we can capture individual cell dynamics in detail. Furthermore, it is an extension of the *NAStJA* framework [24]. Being specifically designed for deployment in High Performance Computing environments, its efficient scaling behaviour allows the simulations of even macroscopic tissues [17, 25]. Hence, CiS has already been used for simulating tissues composing millions of cells [17]. Here, we briefly outline the main properties of the individual layers. A detailed description can be found in [17].

#### CPM layer

The CPM is an extension of the Potts model, developed by Graner and Glazier [26]. In it, we describe cells as connected voxels of same value on a three-dimensional regular grid. The value of individual voxels may change over time depending on the energy of the system. This energy is described by the following Hamiltonian:

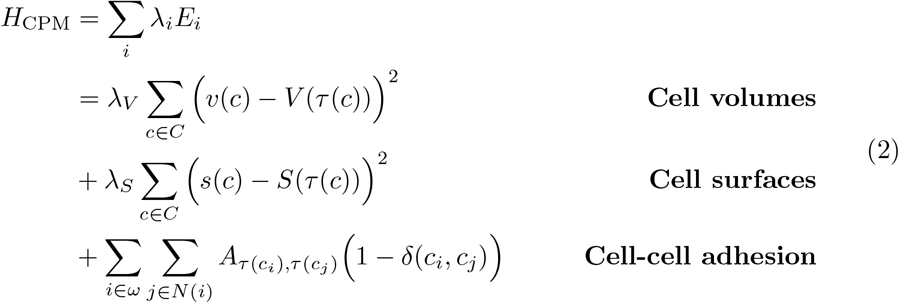

where *λ*_*i*_ is the weight factor for the energy contribution *E*_*i*_, *c* is a cell from the set of all cells *C, τ*(*c*) is the type of cell *c, s*(*c*) and *v*(*c*) are the current surface and volume of cell *c, S*(*τ*) and *V* (*τ*) are the target surface and volumes of cells of type *τ, A* is the adhesion coefficient matrix for all cell types, *N*(*i*) are all lattice points neighboring point *i*, and *δ* is the Kronecker delta. Equation 2 can be extended to include further effects, such as cell motility [27]. In order to propagate the system, we utilize the Metropolis algorithm [28]. First, we change the values of randomly picked voxels into those of a respective neighboring voxel and calculate the energy difference Δ*E* between the old and new system state:

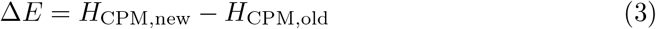

We then decide whether to accept or reject the change. The acceptance probability *p*_accept_ is calculated as follows:

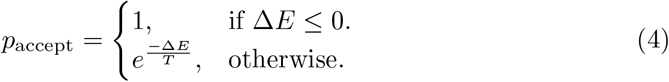

Here, *T* is the temperature of the system. Upon successive propagation steps, cells will expand, compress, deform and move, thereby mimicking their behavior in real tissue. CiS also includes the possibility to add “solid” voxels to the system. Such voxels cannot change their value, and no other voxels can assume a value associated with a solid. This enables us to add structures such as fixed blood vessels.

#### Compound-exchange layer

In addition to detailed cellular structure, CiS contains functionalities for adding compounds such as signals or nutrients to the system. These are exchanged between cells via the cell-cell interface. Solids on the CPM layer can act as sources, thereby mimicking nutrient supplying blood vessels. The flux 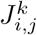 of compound *k* between two cells *i* and *j* is calculated as:

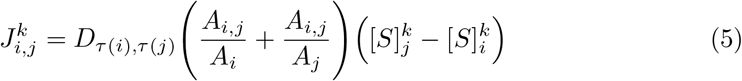

where *D*_*τ*(*i*),*τ*(*j*)_ is the diffusion constant between cells of type *τ*(*i*) and *τ*(*j*), *A*_*i,j*_ is the interface area between cell *i* and cell *j, A*_*i*_ and *A*_*j*_ are the overall surface areas of cell *i* and cell *j*, and [*S*]_*i*_ and [*S*]_*j*_ are the compound concentrations within each cell. The aforementioned solids in the CPM layer can also function as compound sources or sinks. The impact of the compound on the system is defined on the agent based layer.

#### Agent based layer (ABL)

On the ABL, each cell within the tissue is tracked as an individual agent. The properties of each agent are adjusted based on information from the other two layers, and their effects are calculated. Furthermore, cellular functions that are not intrinsically captured by the CPM or the diffusion model, such as cell division and cell death, are implemented here. The use of an ABL also allows us to include cell mutation into the framework: upon each cell division, the properties of the two daughter cells can be changed, such that new cell types emerge.

By combining all three layers, we gain a versatile tool, which can then be parameterized to describe a wide range of system dynamics.

### 4.2 Model parameters

CiS contains many parameters which influence the behavior of the simulated tissue. Here we highlight those used in this study.

#### Cell-cell adhesion

Adhesion interactions between cells are known to vary within tissues, and especially within tumors. They mediate tissue fluidity and effects such as the epithelial-to-mesenchymal transition (EMT) demonstrate their importance [29]. In our simulations, healthy cells adhere to each other and to tumor cells at a strength of 50 AU, which we define as the baseline adhesion. Tumor cells can have an adhesion strength between 0 and 140 AU (see *Cell mutation* paragraph below).

#### Cell motility magnitude and persistence

The CPM Hamiltonian of CiS includes the possibility of adding a cell motility term [27]:

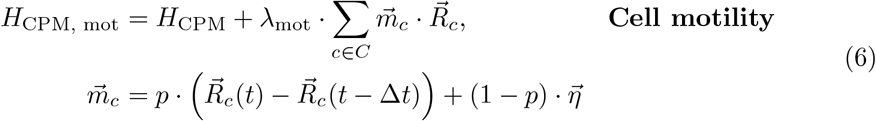

Here 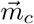 represents the potential and direction of a cell *c, R*_c_(*t*) and *R*_c_(Δ*t*) represent the center of mass of cell *c* at MC step *t* and Δ*t*, respectively, and 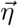 is a random vector obtained by a Wiener process [30]. Each cell follows a modified persistent random walk. The energy contribution of each cell is the dot product of 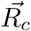 and 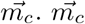 is obtained using the cell’s previous movement and the random vector 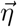. By varying the persistence parameter *p* ∈ [0, 1], we can mediate the contribution of persistent and random movement. At *p* = 0 a cell performs a purely random walk, while at *p* = 1 it performs a purely persistent walk [31]. Finally, we mediate the coupling strength of the motility term to the CPM via *λ*_mot_. In our simulations, healthy cells have a motility strength of *λ*_mot,healthy_ = 0 AU. The motility of tumor cells varies between 0 and 105 AU (see *Cell mutation* paragraph below).

#### Nutrient content

Cells require nutrients to proliferate, and a growing tumor is strongly limited by the nutrients supplied by the surrounding vasculature. We therefore defined a “nutrient” compound on the diffusion layer. For simplicity, this single nutrient combines oxygen and all other nutrients supplied by blood vessels. It is released from solids on the CPM layer to adjacent cells. The solids act as a reservoir with a constant concentration of 4 AU. Therefore, the nutrient amount of all cells lies between 0 and 4 AU. The diffusion constant between vessels and cells, and between any two cells is *D* = 0.1 AU, and diffusion takes place once every 10 MC steps. Finally, each cell consumes nutrients: healthy cells use 0.05 AU nutrients every 10 MC steps. We assume that tumor cells are less efficient and therefore use 0.1 AU nutrients every 10 MC steps [32].

#### Cell division rate

Cell agents within CiS are capable of cell division. When this occurs, the cell is split across a randomly oriented division plane on the CPM layer. Half of the voxels belonging to the old cell are assigned a new value which corresponds to the ID of the new cell. The volume, surface, age and generation properties of the old cell agent are then adjusted, until finally both cells resemble daughter cells of the original one. In order for division to occur, a division condition must be fulfilled. This condition is customizable and can include a number of different terms, e.g. oxygen content, cell volume, cell generation and a random component. Since the healthy tissue in our simulations are meant to model a coarse-grained tumor environment, only the tumor cells are capable of division. Our assumption here is that the surrounding tissue is in a steady-state of division and death. In order for a tumor cell to divide, its volume must be at least 60 % of its target volume, and its nutrient content must be greater than or equal to 1 AU. Furthermore, we set the division probability such that a cell divides roughly every 10 000 MC steps.

#### Cell mutation

Upon division, a cell can mutate, thereby changing some of its agent properties. We decided to focus on two commonly used properties: the cell motility and adhesion strength [33]. A cell has a 10 % chance to increase or decrease either its motility or adhesion strength upon division. Since we did not see significant influence of the mutation properties on the simulations, we only briefly mention this here.

#### Cell death

While cell agents are capable of division, they can also die. In real tissues cells can die due to multiple reasons, but for simplicity here we decided to only focus on the nutrient availability. Hence, a cell will die if its nutrients drop below 0.1 AU.

### 4.3 Loading of blood vessel data into CiS

CiS is parallelized using the Message Passing Interface (MPI), and therefore workers have sperate memory address spaces [17]. Each worker is used to simulate a sub part of the overall CPM volume, which we call a block. We have implemented the loading of the blood vessel data such that each worker loads its individual file at the start of the simulation. These files are generated prior to running the simulation. We first specify the number and size of blocks in each dimension, and split the blood vessel binary mask into accordingly sized sub volumes. These sub volumes are then written out as files named *n*.*raw*, with *n* being the MPI rank to which the sub volume belongs. Each file contains a list of voxel values, which is the flattened version of the respective sub volume. The flattening is done in C-order (row major).

### 4.4 Data visualization

The visualization of the three-dimensional binary masks and the simulations presented a challenge, since non-specialized data visualization frameworks have difficulty to do so at this scale (which are up to giga or even tera-voxel). It was achieved using volcanite (to be published) based on previous work [34].

## Acknowledgements

The authors gratefully acknowledge the Gauss Centre for Supercomputing e.V. (www.gausscentre.eu) for funding this project by providing computing time through the John von Neumann Institute for Computing (NIC) on the GCS Supercomputer JUWELS at Jülich Super-computing Centre (JSC). We thank Max Piochowiak for insightful discussions. We are grateful for funding by the Helmholtz Association in the programs Helmholtz AI (AS, JH) and HIDSS4Health (AS, JK). The funders had no role in study design, data collection and analysis, decision to publish, or preparation of the manuscript.

## Code and data availability

The code of CiS can be found at https://gitlab.com/nastja/nastja. The data and analysis scripts required for reproducing our analysis can be found at https://gitlab.jsc.fz-juelich.de/behle2/mousebrainvasculaturepaper. Due to the size of the entire simulation data (roughly 1.1 TB), we only include the last output file of each simulation within this repository. The full data are available upon reasonable request.

## Declarations

The authors declare that they have no competing interests.

## Notes

https://gitlab.com/nastja/nastja

### Competing Interest Statement

The authors have declared no competing interest.

