## SupplementalInformation for "Leveraging Experimental Vasculature Data for High Resolution Brain Tumor Simulations"

**Supplementary information.**

### Appendix A Mouse brain data processing: stitching multiple stacks

In order to fully utilize the data and not be limited by the geometry of a single stack, we implemented a stitching procedure. This procedure combines parts of multiple microscopy stacks into a composite volume. For this, first the desired volume and location within the raw data is defined. Next, all microscopy stacks that have at least partial overlap with the specified volume are determined. Then, the overlapping part of each binary mask is extracted and added to the stitched volume. While doing this, care must be taken to include the overlap of 300 µm between adjacent stacks specified by Di Giovanna et al. However, even doing so is not sufficient: we still noticed some residual inconsistency at the interface between stacks (see Figure S1). To alleviate this, we varied the overlaps in x- and y-direction within specified bounds and calculated the agreement $a$ between the overlapping parts of both masks for each variation. $a$ was calculated by applying an XNOR between each voxel of the overlap, summing the results and dividing by the number of voxels. Finally, we chose the variation with the highest value of $a$. While there are remaining artefacts inherent to the measurement procedure in Figure S1, this has led to some improvement. While we did not use the stitching procedure for simulations in this study, it represents a useful way of building compound structures.

### Appendix B Supplementary tables

**Table B1 Simulation parameters.**

| Parameter | Value |
|---|---|
| Number of different vessel structures | 10 |
| Number of simulations per vessel structure | 8 |
| Cell target volume | 5000 |
| Cell target surface | 1856 |
| Cell-cell adhesion | 20, 140 |
| Cell motility magnitude | 0-105 |
| Cell-vessel adhesion | 20-140 |
| Random walk persistence | 0.2 |
| Vessel nutrient content | 4 |
| Cell division probability per MC step | 0.0001 |
| Cell division minimum nutrients | 1.0 |
| Cell death nutrient threshold | 0.1 |

Listed are either fixed parameters or ranges, if the respective parameter was varied, e.g. through cell mutation.

**Table B2** **Mouse brain data analysis parameters.**

| Parameter | Value |
|---|---|
| Number of microscopy image z-stacks | 154 |
| Number of microscopy images per z-stack | 2160 |
| Microscopy image resolution | 2048 x 2048 px |
| Microscopy image area per pixel | 0.65 x 0.65μm$^2$ |
| Binary mask building threshold filter value | 175 |
| Stitching procedure maximum alignment shift | 50 |
| Stitching procedure stepsize | 2 |
| Fit values for main manuscript equation 1 | $A \approx 1.877 \cdot 10^5, B \approx 31.77, C \approx 262.9$ |

Listed are the parameters related to the mouse brain data processing pipeline. Raw data generated by Di Giovanna et al. [1].

### Appendix C Supplementary figures

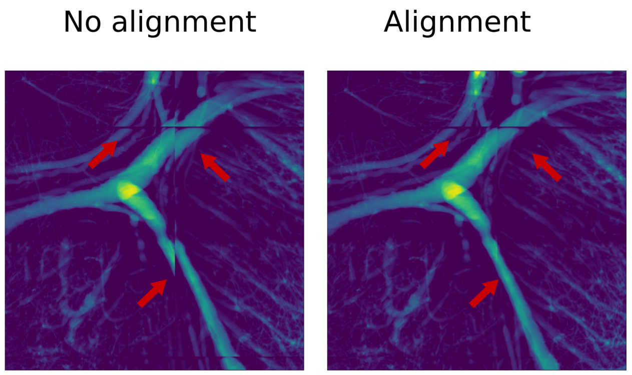


**Fig. S1** SI: Result of custom alignment in microscopy stack stitching procedure. Left: no additional alignment, only the overlap of 300 μm specified in [1] is included. This results in imperfect alignment of some blood vessels (see red arrows). Right: custom alignment included. Adjacent vessels match up much better.

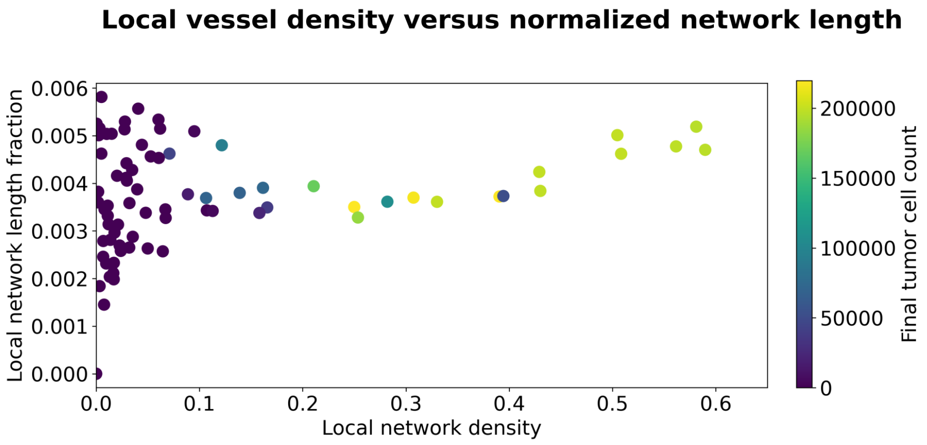


**Fig. S2** SI: Mousebrain vessel property comparison. Shown are the local blood vessel density versus the local blood vessel network length for eight simulations of each respective microscopy stack chosen in main manuscript section 2.1.4. In each simulation, the tumor was placed at a different starting position, and hence the local vessel properties differ. The color indicates the final tumor cell count of the respective simulation.

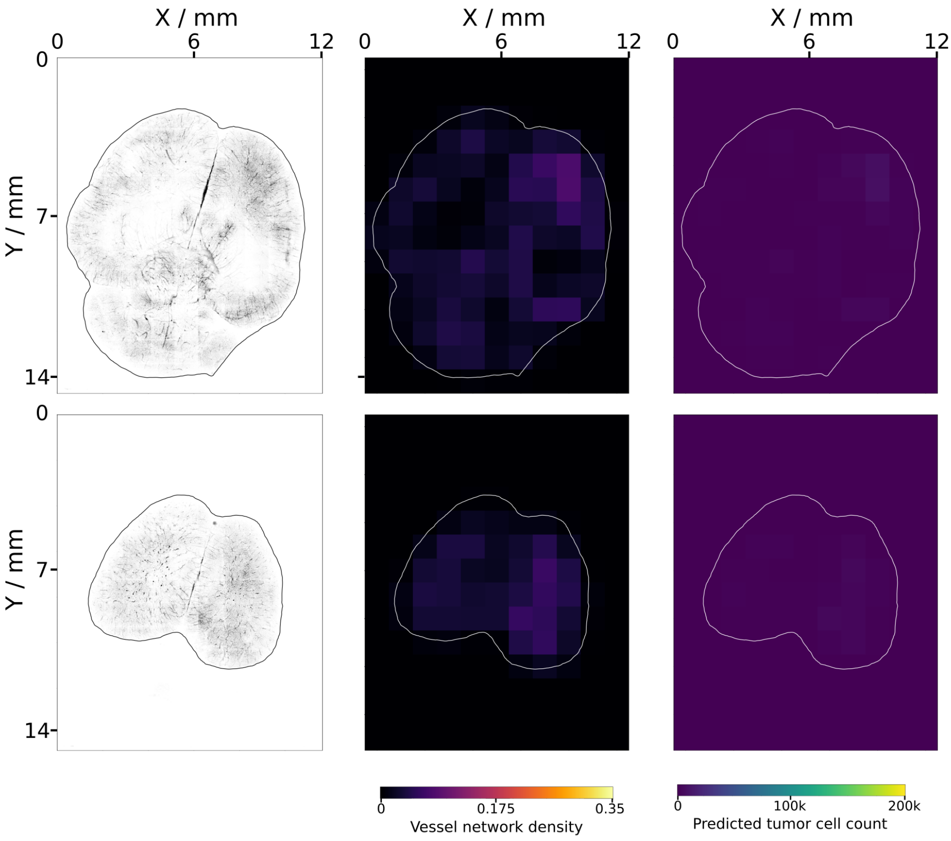


**Fig. S3** SI: Mouse brain blood vessel density and predicted tumor growth. Left: Top-down views of the processed and stitched microscopy stacks for the upper two layers of the brain. Middle: heatmaps showing the blood vessel volume fraction for each microscopy stack. Right: heatmaps showing the predicted tumor cell count for each microscopy stack. Predicted values were obtained using main manusript equation 1.

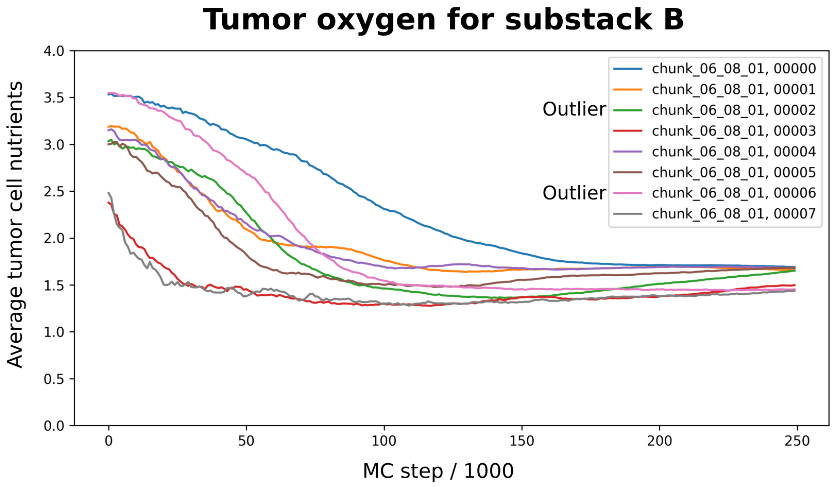


**Fig. S4** SI: Tumor cell oxygen content over time for the simulations within the vasculature of substack B (see main manuscript Figure 3). Simulation 2 and 6, which represent the two outliers found in main manuscript Figure 5 b), both start at high nutrient content, with the nutrients of 6 being higher. However, due to a difference in starting position, over the course of the simulation the environment of simulation 2 outside of the sphere in which the local density was calculated is more favourable, and the subsequently the tumor grows faster. Both are hindered by their overall environment and therefore do not reach a final cell count comparable to 0, 1, 4 and 5

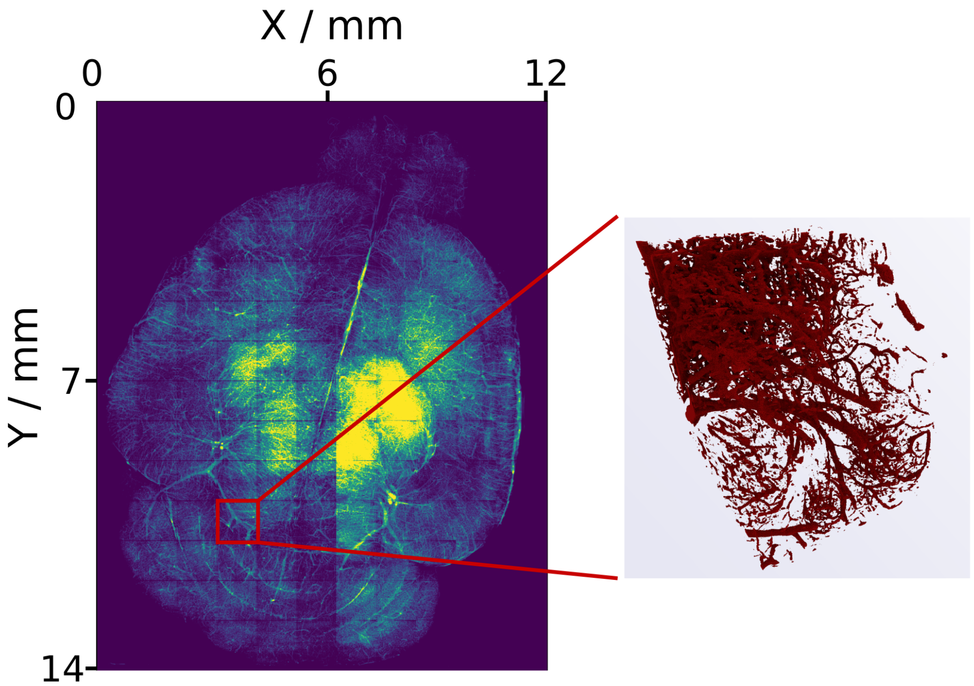


**Fig. S5** Mouse brain vasculature data overview. Left: Top-down view of the entire mouse brain. This image consists of 154 concatenated microscopy stacks, indicated by the rectangular artifacts. Each microscopy stack contains 2160 images. Right: 3D-rendering of a single processed microscopy stack. Raw data produced by Di Giovanna et al [1].
